# IGFs regulate cancer cell immune evasion in prostate cancer

**DOI:** 10.1101/2024.12.03.626600

**Authors:** Ashwin M. Nandakumar, Alessandro Barberis, Jinseon Kim, Cameron R. Lang, Jack V. Mills, Guillaume Rieunier, Dimitrios Doultsinos, Avigail Taylor, Ashwin Jainarayanan, Su M. Phyu, Leticia Campo, Alistair Easton, Eileen Parkes, Timothy James, Freddie C. Hamdy, Clare Verrill, Ian G. Mills, Valentine M. Macaulay

**Affiliations:** Nuffield Department of Surgical Sciences, University of Oxford, Oxford, UK; Department of Oncology, University of Oxford, Oxford, UK; Kennedy Institute of Rheumatology, Nuffield Department of Orthopedics, Rheumatology and Musculoskeletal Sciences, The University of Oxford, Oxford, UK; Department of Biochemistry, Oxford University Hospitals NHS Foundation Trust, Oxford, UK; Department of Cellular Pathology, Oxford University Hospitals NHS Foundation Trust, Oxford, UK; Oxford Cancer Centre, Churchill Hospital, Oxford, UK

**Keywords:** Prostate cancer, IGF-1, immune, PD-L1, antigen presentation

## Abstract

Insulin-like growth factor-1 (IGF-1) promotes prostate cancer (PCa) development and lethality, with immunosuppressive properties in other disease models. These studies investigated the tumor-intrinsic immune effects of IGF-1 in PCa to understand mechanisms underlying its poor immunotherapy response. Transcriptional profiling of human (DU145, 22Rv1) and murine (Myc-CaP) PCa cells revealed, through pathway enrichment, that cytokine signalling, antigen processing and presentation, and other immune regulatory pathways, were most suppressed by IGF-1. We went on to investigate changes in the expression of components of two key pathways responsible for cancer cell recognition by immune cells and immune evasion: antigen processing and presentation, and PD-L1 checkpoint expression. These pathways are crucial for determining immunotherapy response. IGF-1 downregulated transporters associated with antigen processing (*TAPs*), endoplasmic reticulum aminopeptidase-1 (ERAP-1), and Class I component β2-microglobulin, without major changes in Class I allele expression. These effects were associated with reduced presentation of Class I complexes on the Myc-CaP cell surface suggesting altered peptide transport, processing, and/or presentation. In contrast, IGF-1 upregulated immune checkpoint *CD274* (PD-L1) via IGF receptor/AKT/ERK-dependent signaling. Analysis of public data (TCGA Firehose Legacy PCa) revealed increased *CD274* expression in PCa with high endogenous *IGF1* and *IGFBP5*, markers of high IGF axis activity. Additionally, in primary PCa (n=32), multiplex immunofluorescence revealed higher PD-L1 expression in the central tumor of men with high serum IGF-1 promoting cancer cell immune evasion in PCa. These findings indicate a novel mechanism by which IGF-1 mediates immunosuppressive effects in malignant prostate epithelium, potentially explaining the poor responsiveness of PCa to immunotherapy and highlighting the complex interplay between IGF signaling and immune evasion.

## Introduction

Insulin-like growth factors (IGFs) promote normal development and contribute to the growth and spread of cancers, primarily mediated by activation of type 1 IGF receptors (IGF-1Rs) (1). We previously showed that IGF-1Rs are overexpressed in primary prostate cancer (PCa) and are further upregulated on progression to androgen-independent metastatic disease (2, 3). IGF-1 also influences cancer risk: men with high serum IGF-1 have increased risk of PCa development and lethality, with evidence of a causative association (4, 5). Although many pro-tumorigenic effects of IGF axis activation are mediated via IGF-1Rs expressed by target epithelial, mesenchymal, and endothelial cells, it is increasingly recognized that IGFs also influence anti-tumor immunity. IGF-1 promotes the function of immunosuppressive FOXP3+ regulatory T-cells (Tregs) and enhances T-cell secretion of cytokines including IL10, required for macrophage polarization to the pro-tumorigenic, immunosuppressive M2 phenotype (6–8). These factors may contribute to the strongly immunosuppressive tumor microenvironment (TME) of PCa and poor response to immune checkpoint blockade (ICB) (9).

Cancers can evade detection by the adaptive immune system by suppressing cell surface presentation of antigenic peptides. Generally, aberrant cells are recognized by the presence of short peptides generated in the proteasome, transported into the endoplasmic reticulum (ER) by transporters associated with antigen processing (TAP1/2), trimmed by ER aminopeptidases (ERAPs) to modify affinity for Class I molecules, and loaded onto Class I heterodimers comprising a polymorphic heavy α-chain and an invariant β2-microglobulin (β2M) light chain for presentation at the cell surface (10, 11). Many cancers including PCa downregulate cell surface Class I (12); it was shown 30 years ago by Trojan and colleagues that this defect could be rescued in rat high grade glioma (HGG) cells by IGF-1 depletion, rendering the cells non-tumorigenic and able to induce CD8+ cytotoxic T-cell infiltration and regression of unmodified tumors in syngeneic animals (13). Similar immune-mediated tumor regression was reported in rats injected with IGF-1R-depeted HGG cells, findings that triggered a pilot clinical trial and a Phase IIb trial (NCT04485949) (14–16).

Clinical evidence for IGF-associated immunosuppression comes from reports of low tumor infiltrating lymphocytes (TILs) in high IGF-1R PCa bone metastases, and genomic aberrations in cancer patients with hyper-progressive disease post-ICB, including mutations in negative IGF regulator *IGFBP2* and transcriptional upregulation of IGF-1, PI3K-AKT and ERK pathways (17, 18). More recently, short-term starvation was shown to sensitize non-small cell lung cancer (NSCLC) to PD-1 blockade *in vivo* by suppressing serum IGF-1 and IGF-1R activity in tumor tissue. Furthermore, NSCLC patients with durable clinical benefit post-ICB had significantly lower circulating IGF-1 and lower tumor IGF-1R than those with no durable benefit (19). IGF-1R inhibition is also reported to enhance responses to anti-PD-1 in colorectal cancer and triple negative breast cancer (20, 21). These findings prompted us to investigate the hypothesis that IGF-1 has immunosuppressive actions in prostate cancer epithelium contributing to cancer cell immune evasion.

## Methods

### Cells, reagents

Human PCa cell lines DU145 and 22Rv1 were from Cancer Research UK Laboratories (Clare Hall Hertfordshire UK) and Professor Sir Walter Bodmer (University of Oxford, UK) respectively. Cell lines were cryopreserved at early passage, regularly Mycoplasma tested and used within 20 passages. BMS-754807, AZD5363 and trametinib were from Selleck Chemicals.

### Reverse-transcription, quantitative PCR (qPCR)

RNA was extracted with the RNeasy kit (Qiagen) and reverse transcribed with SuperScript III First-Strand Synthesis SuperMix (Invitrogen). Gene expression was measured on the 7500 Fast Real-Time qPCR System (Thermo Fisher) using SYBR Green (New England Biolabs) and primers listed in Supplementary Table S1.

### Flow cytometry

Cultured cells were dissociated with 2 mM Ethylenediamine tetra-acetic acid (EDTA, Sigma) in PBS at 37 °C, collected and washed in cold MACs buffer (0.5% bovine serum albumin, BSA, 2 mM EDTA in PBS). Cells were stained using antibodies listed in Supplementary Table S2 diluted in MACs buffer, then washed, fixed and stored in the dark at 4°C. Surface expression was measured on the Attune NxT Flow Cytometer, data were analyzed using FlowJo 10.8.1 and expressed as % positivity and median fluorescence intensity (MFI).

### Multiplex Immunofluorescence, HALO analysis

Multiplex immunofluorescence (mIF) was performed on 4 μm FFPE sections of radical prostatectomies from men recruited to the Prostate Cancer Mechanisms of Progression and Treatment (ProMPT) study, as described in (22) and mIF methods in (23), using Epitope Retrieval (ER) solutions, primary antibodies and their opal fluorophore pairings listed in Supplementary Table S2. Benign and cancer areas were annotated by Uro-Pathologist (CV), and areas of epithelium, stroma and artifact identified using the internal MiniNet AI classifier. Tumor margin, cancer and cancer center areas were annotated by HALO, and cells detected and signals quantified on the HALO HighPlex FL v3.1.0 module (Indica Labs).

### RNA-sequencing (RNA-seq)

RNAs were quantified using RiboGreen (Invitrogen) on the FLUOstar OPTIMA plate reader (BMG Labtech) and size profile and integrity analyzed on the 2200 or 4200 TapeStation (Agilent, RNA ScreenTape). Input material was normalized to 20 ng, ribosomal RNA depleted using NEBNext rRNA Depletion Kit (NEB, Human/Mouse/Rat) and strand-specific library preparation used NEBNext Ultra II mRNA kit (NEB) following manufacturer’s instructions. Libraries were amplified (17 cycles on Tetrad, Bio-Rad) using in-house unique dual indexing primers based on (24). The size profile of individual libraries was analyzed on the 2200 or 4200 TapeStation and normalized using Qubit. Libraries were pooled, diluted to ∼10 nM, denatured, and further diluted for paired-end sequencing (NovaSeq6000 platform, Illumina, NovaSeq 6000 S2/S4 reagent kit v1.5, 300 cycles).

### Statistical and Bioinformatic analysis

Using GraphPad Prism v9, we applied t-tests to calculate significance between two treatment groups, one-way ANOVA with Tukey’s multiple comparison test for three or more treatment groups and Log-rank (Mantel-Cox) test for clinical survival data. Graphs show data from three independent experiments and p values <0.05 were considered statistically significant. RNA-seq data were analyzed using software listed in Supplementary Table S3. The false discovery rate was controlled using the Benjamini-Hochberg (BH) method and results considered significant at adjusted p-value <0.05. Gene set enrichment analysis (GSEA) used gene sets from the Molecular Signatures Database. For immune-focused analysis, we manually selected pathways from Gene Ontology (GO) Biological Processes, KEGG, BioCarta, WikiPathways, Reactome, and Hallmarks MsigDB collections, grouping them into three categories: cytokine, antigen processing/presentation and ‘other immune-related’. Enrichment analysis used the clusterProfiler R package with BH multiple testing correction. Results were considered significant at adjusted p-value <0.05, Venn diagrams and heatmaps were created with EnhancedVolcano, ggVennDiagram and ComplexHeatmap R packages, and GSEA plots with clusterProfiler (see Supplementary Table S3).

## Results

### IGF-enriched pathways include activated cell cycle, DNA replication and repair and suppressed immune pathways

As a first step to investigate associations of IGF axis components with immune factors, we analyzed effects of IGF-1 on gene expression in three PCa cell lines: two human, androgen receptor (AR)-positive 22Rv1 cultured from a xenograft established from primary PCa, and AR-negative DU145 derived from metastatic cancer; and one murine, Myc-CaP, derived from primary adenocarcinoma in the Hi-Myc mouse (25). Sub-confluent cultures were serum-starved for 24 hr and treated with IGF-1 or solvent (control) for 24 hr prior to RNA extraction. We identified 3,471 differentially regulated genes (DEGs) in DU145, 2,321 in 22Rv1 and 600 Ensembl notations corresponding to 584 gene IDs in Myc-CaP (Supplementary Tables S4-6, Supplementary Figure S1). Variation in the numbers and identity of IGF-regulated genes may reflect differences in AR status and potentially the dominance of Myc transcriptional programming in Myc-CaP cells (26). There were 31 significantly upregulated DEGs and 26 downregulated DEGs in common between the 3 cell lines (Figure 1A). These included 5 genes in the COSMIC Cancer Gene Census of mutated genes implicated as cancer drivers (Supplementary Table S7), and 12 upregulated and 10 downregulated genes associated with immune-relevant pathways (Figure 1B).

**Figure 1:**
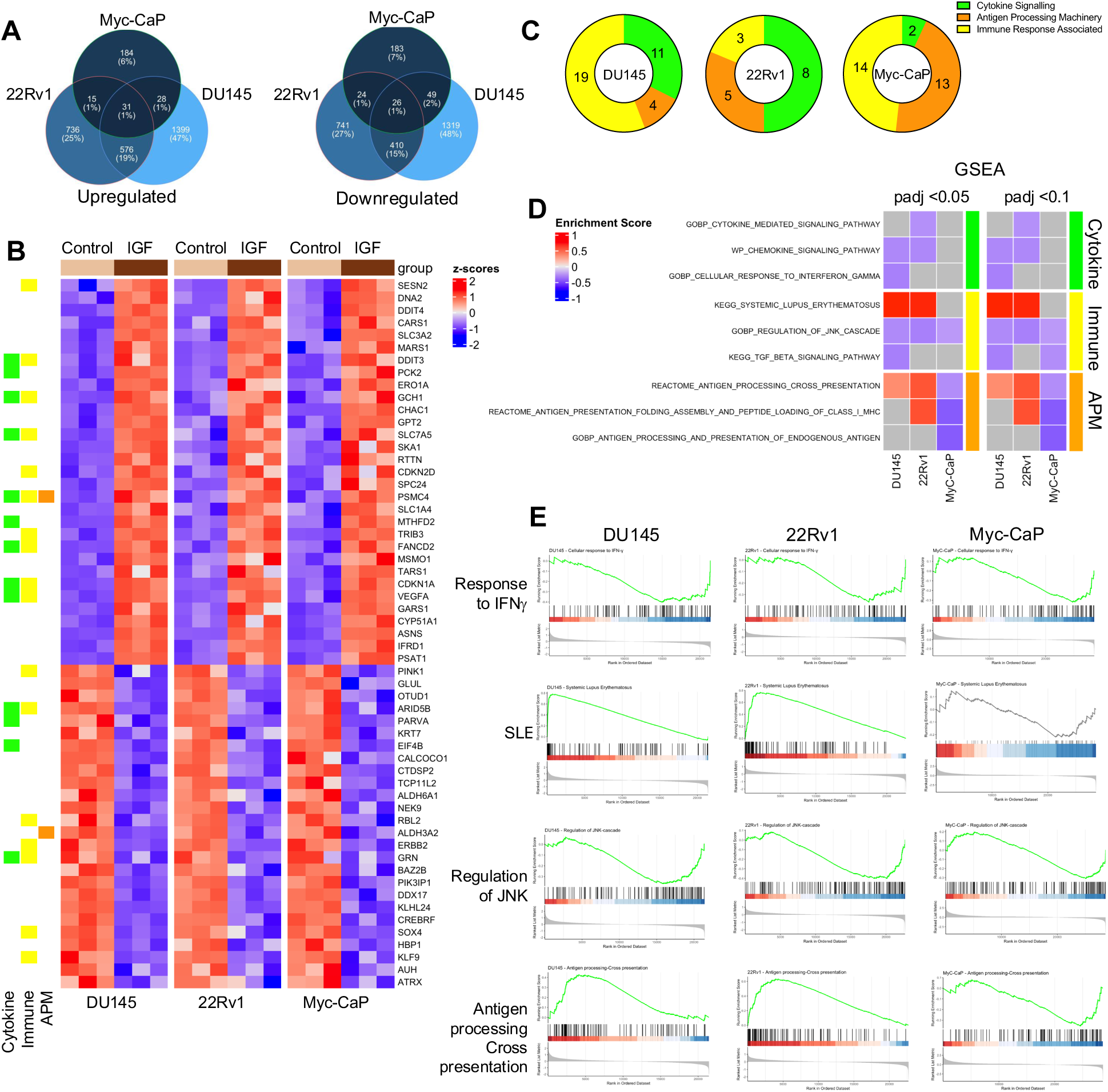
IGF-1 deregulates immune-associated genes and transcriptional pathways. Subconfluent DU145, 22Rv1 and Myc-CaP cultures were serum-starved for 24 hr, stimulated with 30 nM IGF-1 or solvent (control) for 24 hr and RNA was extracted for RNA-seq. **A.** IGF-induced DEGs unique to and shared between the 3 PCa cell lines showing number (%). **B.** Expression heatmap of 57 DEGs in common between the 3 cell lines including 31 significantly upregulated and 26 downregulated DEGs and annotation into three curated immune relevant categories: cytokine signaling, immune response associated and Antigen Processing Machinery (APM). **C.** GSEA was performed using the three curated immune-associated pathway datasets (Supplementary Table S11). Pie charts represent the distribution of enriched pathways in each cell line. **D.** Heatmap of selected enriched pathways and enrichment scores at padj<0.05 and <0.01 for each cell line. **E.** GSEA enrichment plots showing response to interferon gamma (IFNγ), systemic lupus erythematosus (SLE), regulation of JNK cascade and antigen processing and cross presentation.

Initial unbiased enrichment analysis highlighted activation of cell cycle, DNA replication and repair, and metabolism-related pathways (Supplementary Tables S8-10). IGF-induced regulation of these pathways is well documented (1), providing confidence that the cells generated expected responses to IGF-1. Enrichment analysis also identified pathways less well-recognized as IGF-related, including immune regulator-associated pathways, almost all with negative enrichment scores suggesting pathway inactivation (Supplementary Tables S8-10). This finding prompted us to perform focused enrichment analysis on pathways related to three broad categories of immune function (Supplementary Table 11). These were production of and response to cytokines and chemokines, antigen processing machinery (APM), and other immune response-related pathways including checkpoint regulation.

The enrichment pattern varied between cell lines, with predominant association with cytokine signaling pathways in DU145, cytokine signaling and immune response-related in 22Rv1, and antigen processing and presentation in Myc-CaP cells (Figure 1C, Supplementary Table S12). Figure 1D shows a heatmap of enriched pathways and enrichment scores for each cell line. Amongst cytokine-related pathways, the cellular response to interferon gamma (IFNγ) was suppressed in all 3 cell lines (Figure 1E). General ‘immune-related’ pathways included regulation of JNK cascade, inactivated in all 3 cell lines, and ‘systemic lupus erythematosus’, activated in DU145 and 22Rv1 and containing many complement, Fc receptor and HLA encoding genes (www.genome.jp/entry/pathway+hsa05322). There was also enrichment of APM-related pathways, apparently activated in the human cells but suppressed in Myc-CaP (Figure 1D-E).

### IGF-1 influences expression of genes that regulate protein degradation, peptide processing and presentation

Defects in antigen presentation have been described in PCa cell cultures (27), involving multiple components of the APM. Defective APM results in cancer cell immune evasion, diminishing the effectiveness of immunotherapy. Having identified IGF-enriched immune pathways related to antigen processing and presentation (Figure 1D-E) we assessed effects of IGF-1 on expression of genes that regulate intracellular protein degradation, peptide processing and presentation. First we assessed proteolytically active proteasome subunits *MECL1*, *LMP2* and LMP7 required for assembly of the 20S proteasome, protein degradation and amino acid recycling (28), finding IGF-induced *MECL1* downregulation in DU145 cells only and *LMP2* downregulation in both DU145 and 22RV1 (Supplementary Figure S2A). In all three cell lines lysosome-associated membrane protein 2 (*LAMP2*), which maintains lysosomal integrity and function, was downregulated by IGF-1; indeed low *LAMP2* reportedly associates with adverse overall survival in high-grade PCa (29). RB1-inducible coiled-coil protein 1 *(RB1CC1*), an ULK1 complex component required for autophagy induction, was also coordinately downregulated. Autophagy-related genes *ATG14 and VTI1B* are required for autophagosome formation; *ATG14* was downregulated in 22Rv1 and Myc-CaP, and *VTI1B* in 22Rv1 only (Supplementary Figure S2B-C). These data suggest that IGF-1 may influence protein/peptide degradation via lysosomal and/or autophagic pathways.

We then assessed genes encoding regulators of peptide transport, processing and presentation. In all three cell lines IGF-1 caused significant *TAP1*, *TAP2* and *ERAP1* downregulation, accompanied in Myc-CaP by downregulation of TAP binding protein *Tapbp* (Figure 2A-B, Supplementary Figure S3A). Although we identified IGF-induced deregulation of pathways related to antigen presentation (Figure 1D), IGF-1 had little effect on expression of genes encoding MHC Class I α-chains themselves, the only significant effect being apparent *HLA-C* upregulation in 22Rv1 (Supplementary Figure S3B-D). In contrast, β2m was significantly downregulated by IGF-1 in all 3 cell lines (Figure 2C). Given identification of enriched APM pathways in IGF-treated Myc-CaP cells (Figure 1B), we used Myc-CaP to assess potential consequences for antigen presentation. Cell surface Class I complexes were detectable in ∼50% of cells, with marked reduction in both % positivity and MFI after 5 days exposure to IGF-1 (Figure 2D-E). These results suggest that IGF-1 impairs the ability of Myc-CaP cells to present antigen in the context of Class I. This effect appears to correlate with downregulation of genes encoding antigen processing components and β2M rather than downregulation of Class I genes themselves.

**Figure 2:**
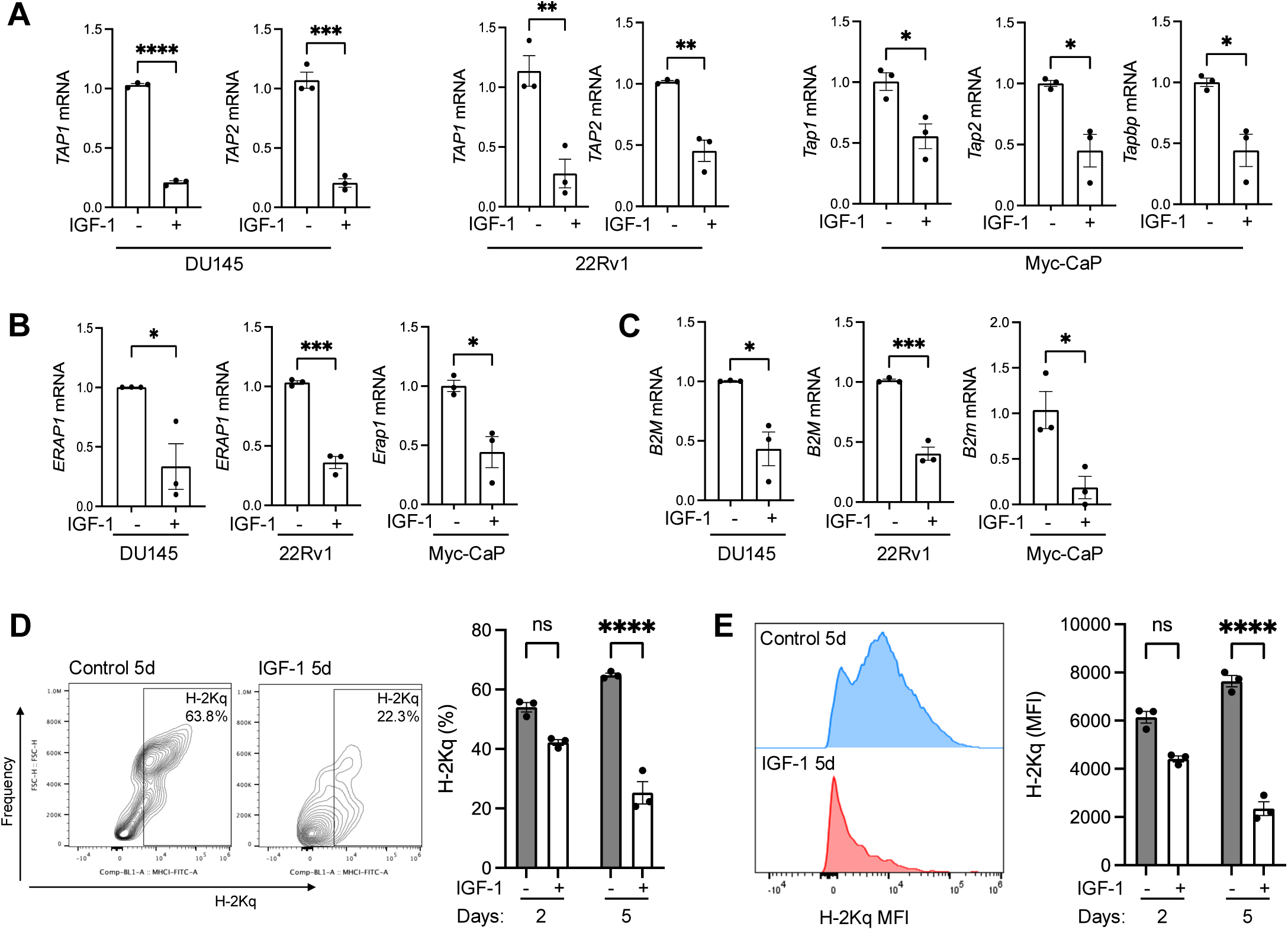
IGF-1 downregulates antigen processing and presentation components. RNA was extracted from DU145, 22Rv1, and Myc-CaP cells 24 hr after serum-starving and IGF-treatment as in Figure 1, and gene expression was assessed by RT-qPCR. Expression of: **A**. *TAP1*, *TAP2* mRNA in all cell lines and *Tapbp* in MycCaP; **B.** ERAP mRNA, **C.** β2M mRNA. **D, E**. Class I complexes are stabilized at the cell surface by high affinity peptide binding, while peptide-free (empty) heterodimers are generally unstable at the cell surface at physiological temperature. We used Class I antibody to H-2Kq, appropriate to the genotype of FVB mice from which Myc-CaP cells were derived (25). Analysis by flow cytometry of H2Kq surface expression in Myc-CaP cells treated with 30 nM IGF-1 or solvent for 2 or 5 days, showing representative flow cytometry plots at 5 days and n=3 independent experiments. D: percentage H2Kq positivity, E: H2Kq MFI. All graphs represent mean ± SEM of three independent analyses (*p<0.05; **p<0.01; ***p<0.001; ****p<0.0001; ns, nonsignificant by one-way ANOVA).

### IGF-axis activation induces PD-L1 upregulation

In addition to a functional APM, the regulation of tumor-intrinsic immune suppressors and immune checkpoint expression are crucial for immune-mediated clearance of cancer cells. *ASS1* and *ARG1* encode enzymes that respectively regenerate and degrade arginine, which has immunosuppressive effects in the PCa microenvironment (30). *ASS1* was downregulated in all three cell lines, *ARG1* in only DU145 (Supplementary Figure 4A-C). There was a non-significant trend to upregulation of *VSIR*, encoding inhibitory checkpoint V-domain Ig suppressor of T cell activation (VISTA), in IGF-treated DU145 and 22Rv1 cells and no detectable *Vsir* in Myc-CaP (Supplementary Figure S4A-B). Other immune checkpoints including *CD274* (PD-L1), *PDCD1LG2* (PD-L2) and *CTLA4* were not identified as IGF-induced DEGs in any cell line, and gene counts for *PDCD1LG2* and *CTLA4* were essentially zero. However, we assessed expression of *CD274* in view of the reported ability of IGF-1R inhibition to synergize with anti-PD-1 immunotherapy (19). In contrast to IGF-induced downregulation of multiple immune-relevant genes detected thus far (Figure 2A-C), IGF-1 caused statistically significant upregulation of *CD274* mRNA in all three PCa cell lines (Figure 3A). By flow cytometry, IGF-1 upregulated PD-L1 surface expression in DU145 cells in both serum-free and serum-supplemented media (Figure 3B, Supplementary Figure S4D). This effect was blocked by pre-treatment with actinomycin D (Figure 3C) consistent with upregulation at the transcriptional level.

**Figure 3:**
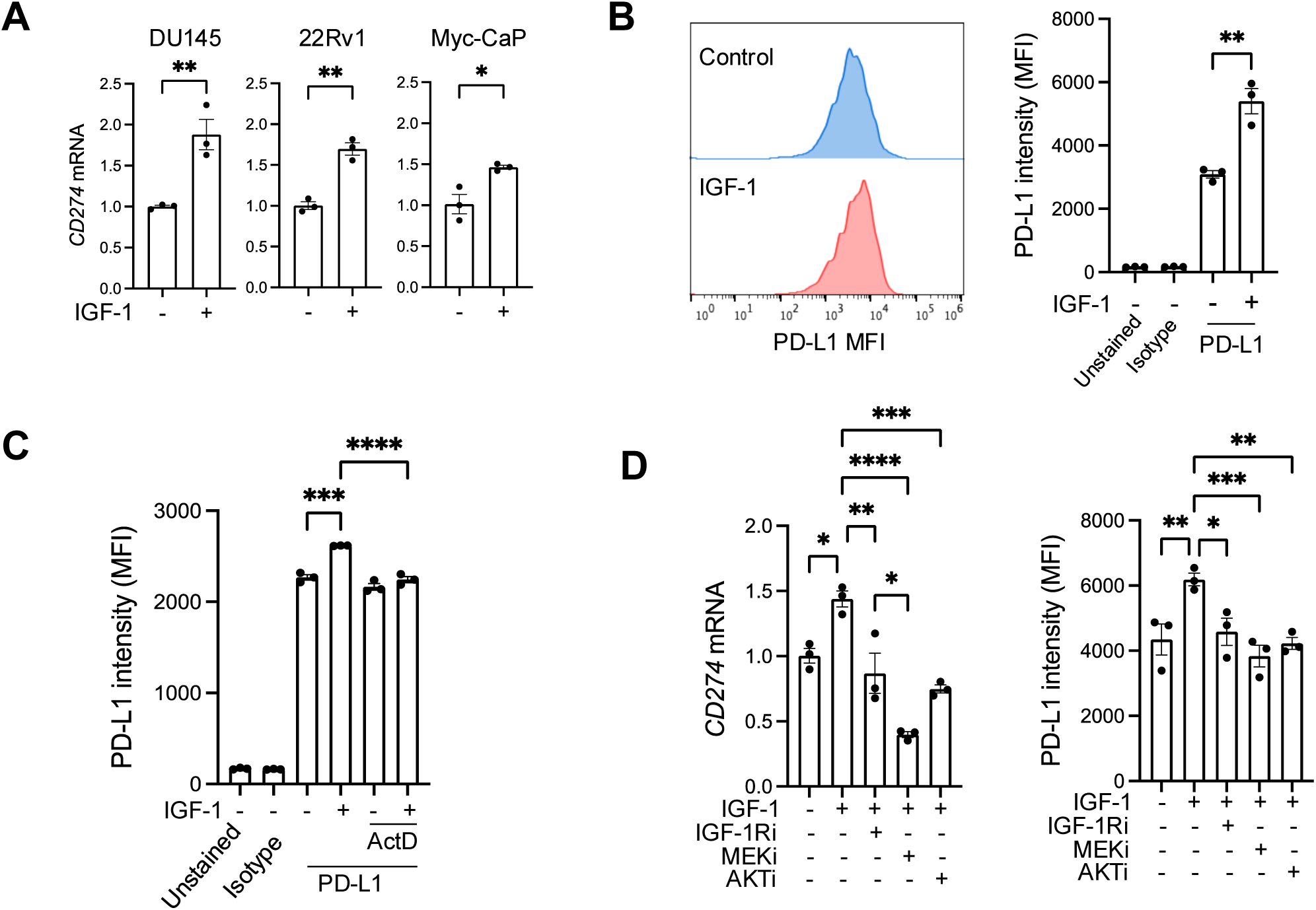
IGF-1 upregulates immune checkpoint PD-L1. **A**. PCa cells were serum starved for 24 hrs, treated with 30 nM IGF-1 or solvent and expression of *CD274* assessed by qPCR (n=3 independent assays, mean ± SEM). **B.** DU145 cells were treated as in A and PD-L1 surface expression measured by flow cytometry. Left, representative flow histogram; right, quantification of MFI (n = 3 independent assays, mean ± SEM). **C.** DU145 cells were pre-treated with 1 μg/mL Actinomycin D for 2 hrs and then 30 nM IGF-1 was added. After 24 hr PD-L1 surface expression was measured using flow cytometry. Graphs represent mean ± SEM. **D.** DU145 cells were pre-treated with 300 nM IGF-1R inhibitor BMS-754807, 5 μM AKT inhibitor AZD5363 and 50 nM MEK inhibitor trametinib for 1 hr and 30 nM IGF-1 was added for 24 hrs. Parallel cultures were used for RNA extraction and RT-qPCR for *CD274* (PD-L1) mRNA quantification (upper graph) and surface protein expression measured by flow cytometry (lower, n=3 independent assays, mean ± SEM). (*p<0.05; **p<0.01; ***p<0.001; ****p<0.0001; ns, nonsignificant by one-way ANOVA).

To investigate how IGF-1 contributes to PD-L1 regulation, we tested effects of inhibiting signaling pathways of IGF-1R and it’s downstream effectors MEK-ERK and AKT. Results suggest that blockade of IGF-1R, MEK-ERK or AKT suppressed IGF-induced PD-L1 upregulation at the transcriptional and cell surface level (Figure 3D). Consistent with our findings, PD-L1 regulation by AKT and MEK was also reported in other human tumor cell lines (31, 32).

### IGF axis activity associates with deregulated immune genes and immune infiltrating cells in PCa tissues

To test the clinical relevance of these findings we interrogated public RNA-seq data from the Prostate Adenocarcinoma Firehose Legacy dataset in cbioportal (www.cbioportal.org/study/summary?id=prad_tcga) (33). First, we tested associations of endogenous *IGF1* expression with *CD274* (PD-L1 mRNA), finding significantly higher *CD274* in tumours with the highest (n=124) quartiles of IGF1 when compared with the lowest (n=125; Figure 4A). As a tissue indicator of IGF axis activation we also assessed IGF binding protein-5 (*IGFBP5*), a well-recognized transcriptional readout of IGF axis activity in many tissues and cell types (34). Indeed, *IGFBP5* expression in this dataset was significantly higher in tumors with high endogenous *IGF1* mRNA, and reciprocally *IGF1* mRNA was higher in high *IGFBP5* tumors (Supplementary Figure S5A,B). Consistent with these associations of *IGF1* and *IGFBP5* mRNA, there was significantly higher *CD274* expression in Firehose Legacy PCa tumors in the highest vs lowest quartiles of *IGFBP5* (Figure 4B). PCa Firehose Legacy tumors expressed low levels of *CTLA4* mRNA, and *CTLA4* transcript levels were significantly higher in tumors expressing high *IGFBP5* although not *IGF1* (Supplementary Figure S5C).

**Figure 4:**
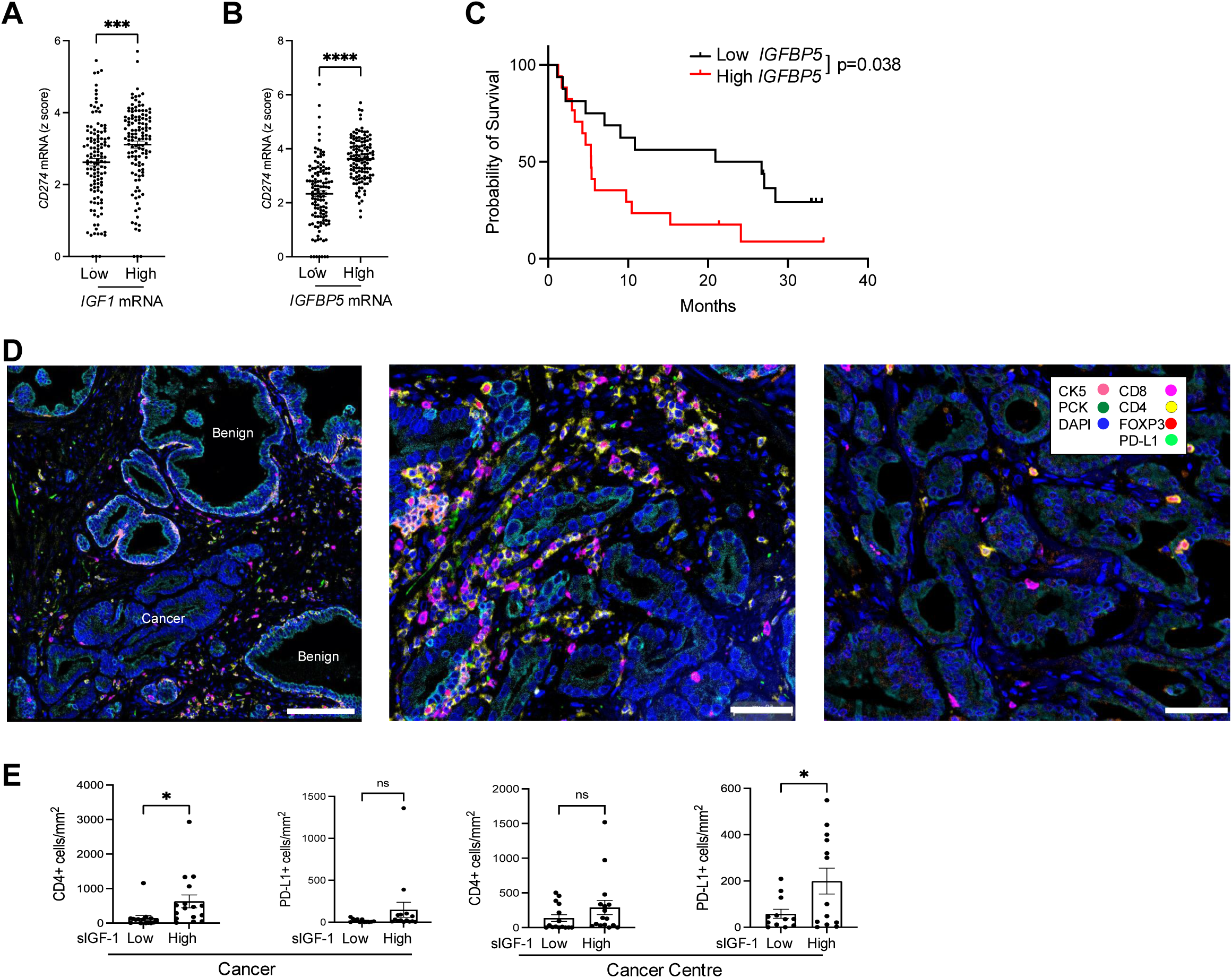
IGF axis markers associate with altered expression of immune genes and immune infiltrating cells in clinical PCa tissue. **A-B**. Tumors from TCGA prostate adenocarcinoma Firehose Legacy dataset (n=498) were divided based on highest and lowest quartiles of tumor endogenous *IGF1* or *IGFBP5* expression (n= low: 124, high =125 in each case). Graphs show *CD274* expression (Z-score relative to all samples) in highest and lowest quartiles of *IGF1* (A) and I*GFBP5* (B). **C.** We accessed data from cbioportal on 110 ipilimumab-treated metastatic melanoma patients, of whom 40 had tumor transcriptional profiling (43). These cases were divided by median *IGFBP5* (not quartiles due to small cohort size). Survival post-ICB censored at 36 months. Overall survival was significantly shorter (Log-rank (Mantel-Cox) test) in patients whose tumors expressed high *IGFBP5* (n=17) vs those with low *IGFBP5* melanomas (n=16). **D.** Multiplex IF of radical prostatectomy tissue of ProMPT cases visualised using HALO. Representative images of: left, junction between benign glands (PCK+ luminal cells surrounded by PCK+/CK5+ basal cells) and malignant epithelium (PCK+/CK5-, scale bar 100 µm); center: high immune infiltrating case; right: low immune infiltrating case (scale bar 50 µm). **E.** Quantification of CD4+ T-cells and PD-L1 positive cells in the total cancer area (left) and cancer center (right) in patients with high or low serum IGF-1. Graphs show mean ± SEM positively-stained cells (n=31 for CD4+ staining, n=25 for PD-L1, remaining cases lost due to technical issues with staining and visualization). (*p<0.05; ***p<0.001; ****p<0.0001; ns, nonsignificant).

We searched for TCGA datasets that included transcriptional analysis and clinical survival data following ICB. We could not identify any datasets including anti-PD-1/PD-L1 therapy, but in a cohort of 110 patients with metastatic melanoma with survival data post anti-CTLA-4 ipilimumab, transcriptional profiles were available in 40. Patients whose tumors expressed high *IGFBP5* had significantly reduced overall survival (Figure 4C). This is consistent with the reported contribution of the IGF axis to ICB resistance (19), although re-testing in larger cohorts will be required to confirm this finding.

As an additional approach to explore associations between the IGF axis and checkpoint expression, we returned to a cohort (n=139) of men with localized PCa (n=139) recruited between 2010-2014 to Prostate Cancer: Mechanisms of Progression and Treatment (ProMPT study, MREC 01/4/061, PI F. Hamdy). We previously identified a transcriptional role for nuclear IGF-1R and association with advanced clinical stage (22). Here, we assayed pre-prostatectomy serum IGF-1, finding that all values were within the normal range for adult men (Supplementary Figure S6A). We selected the 16 cases with the highest (21.74 – 31.49 nmol/L) and 16 with the lowest (7.19 – 11.98 nmol/L) IGF-1 and used FFPE prostatectomy sections for mIF. The mIF panel comprised pancytokeratin (PCK) to stain prostate epithelium, cytokeratin 5 (CK5, expressed by basal cells of benign glands), and markers of helper T-cells (CD4+), cytotoxic T-cells (CD8+), regulatory T-cells (Tregs, FOXP3+) and PD-L1 (Figure 4D). We first identified benign, tumor and central tumor regions of each tissue (Supplementary Figure S6B) and quantified immune and PD-L1 positive cells in each region. Many tumors had sparse immune infiltrates and there were no significant differences in total numbers of infiltrating CD8+ T-cells or FOXP3+ Tregs in the total tumor area or central tumor in high vs low serum IGF-1 cases (Supplementary Figure S6C). However, central tumor (including epithelial and stromal components) of men with high serum IGF-1 contained more PD-L1 positive cells than tumors of men with low serum IGF-1 (Figure 4E, Supplementary Figure S6C). This association is consistent with IGF-induced PD-L1 upregulation in cultured PCa cells (Figure 3). In the total tumor area, there were also more T-cells positive for membrane CD4 in high serum IGF-1 cases compared with low IGF-1 cases (Figure 4E).

## Discussion

Our data indicate that IGF-1 regulates PCa expression of immune associated genes, components of the APM and the immune checkpoint PD-L1, with evidence of enriched and largely suppressed immune pathways. IGF-1 caused consistent downregulation of *TAP1, TAP2*, *ERAP1 and β2M*, suggesting perturbation of the APM. This effect was accompanied in Myc-CaP by TAP binding protein (*Tapbp*) downregulation. These data extend findings reported in HGG cells, where IGF-1 depletion rescued from downregulation of *TAP1*, *TAP2* and *LMP7* (35). Furthermore, we identified significant IGF-induced reduction in presentation of cell surface Class I complexes by Myc-CaP cells. The lack of consistent changes in expression of Class I alleles in human or murine PCa cells suggests that the Class I upregulation and consequent CD8-dependent immune response reported in IGF-depleted or inhibited tumors (13, 19) may be due not to reduced expression of MHC α-chains, but rather to disruption of peptide transport and processing, and downregulation of invariant Class I component β2M. In addition, *ERAP1* downregulation could influence trimming of Class I-bound precursor peptides, which in turn could affect the peptide repertoire and the conformational stability of Class I:peptide complexes (11). As far as we are aware these changes have not been reported in PCa cells, although IGF-1 and insulin are reported to downregulate MHC Class I alleles in FRTL-5 rat thyroid cells and human hair follicle (36, 37). This suggests possible differences in Class I regulation between malignant and non-malignant cells and tissues.

The second concordant change related to *CD274*/PD-L1, reportedly expressed at low levels in primary PCa, with <10% tumors containing IHC-detectable membrane PD-L1 in ≥1% of total cells in malignant epithelia and immune cells (38). Consistent with this, there was evidence of generally low *CD274* mRNA in public PCa data although *CD274* expression was significantly higher in high *IGF1/IGFBP5* tumors (Figure 4A-B) and at the protein level (mIF) in central tumor of men with high serum IGF-1 (Figure 4E).

While IGFs are reported to have both pro– and anti-inflammatory actions, most reports favor an anti-inflammatory role. IGFs suppress cytokine secretion with anti-inflammatory associations in cardiac disease and diabetes (39, 40). Igf-1r activation is reported to suppress inflammation in the TME of NSCLC models (41) and induce degradation of retinoic acid-inducible gene-I (RIG1), a pattern-recognition receptor required for type 1 IFN production and the proper function of the innate immune response (42). Several of our findings are consistent with an anti-inflammatory role in PCa. While single cell studies in IGF-manipulated tissue would be required to assess causality, these changes could suggest that IGF-induced transcriptional deregulation in PCa epithelium induces immunosuppression in components of the PCa TME.

## Acknowledgements

We dedicate this article to the memory of Prof. Valentine Macaulay, our group leader and co-lead author. Prof. Macaulay was not only a brilliant and dedicated scientist but also a compassionate mentor, an inspiring leader, and a cherished friend. Her unwavering commitment to science and her genuine care for those she worked with has left a profound mark on all of us. We feel fortunate to have had the privilege of calling her our colleague.

This study was supported by Cancer Research UK Early Detection project grant (C476/A27060), Prostate Cancer UK awards (RIA-ST2-024 and MA-CT20-006), The Rosetrees Trust and John Black Foundation Charitable Trust (PhD-Plus award PhD2021-100025), Oxford University Human Immune Discovery Initiative (HIDI) fund, and UCARE-Oxford (GR280519). We gratefully acknowledge the contribution to this study made by the Oxford Centre for Histopathology Research and the Oxford Radcliffe Biobank, which are funded by the University of Oxford, the Oxford CRUK Cancer centre, the NIHR Oxford Biomedical Research Centre (BRC) (Molecular Diagnostics Theme/Multimodal Pathology Subtheme and the NIHR CRN Thames Valley network. The ProMPT study was supported by the UK NIHR, Cancer Research UK and the MRC, and the Cambridge and Oxford Biomedical Research Centres. The funding source had no role in the design, conduct of the study, collection, management, analysis and interpretation or preparation, review, or approval of the manuscript.

