## Supplementary Figures for "IGFs regulate cancer cell immune evasion in prostate cancer"

**Nandakumar et al Supplementary Tables and Figures**

**Supplementary Table S7: in this file. The remaining Tables are provided in the Supplementary Excel file:**

**Supplementary Table S1: Primer sequences**

**Supplementary Table S2: Antibodies used for flow cytometry and mIF.**

**Supplementary Table S3: Software availability**.

EdgeR (1)

DESeq2 (2)

GSEA (3)

GO (4)

KEGG (5)

Biocarta (6)

WikiPathways (7)

Reactome (8)

MSigDB (9)

clusterProfiler 4.0 (10)

Complex heatmap (11)

ggVennDiagram (12)

EnhancedVolcano (13)

**Supplementary Table S4: IGF-regulated genes in DU145 cells.**

**Supplementary Table S5: IGF-regulated genes in 22Rv1 cells.**

**Supplementary Table S6: IGF-regulated genes in Myc-CaP cells.**

**Supplementary Table S7: COSMIC genes coordinately deregulated by IGF-1.**

**Supplementary Table S8. Unbiased pathway enrichment analysis by Functional Class Scoring in DU145 cells.**

**Supplementary Table S9. Unbiased pathway enrichment analysis by Functional Class Scoring in 22Rv1 cells.**

**Supplementary Table S10. Unbiased pathway enrichment analysis by Functional Class Scoring in Myc-CaP cells.**

**Supplementary Table S11: Immune-relevant pathways subject to GSEA in cell lines.**

**Supplementary Table S12: Immune-relevant pathways enriched in PCa cells.**

| **Symbol** | **Name** | **Role in cancer** | **+IGF-1** |
| --- | --- | --- | --- |
| ATRX | ATRX, chromatin remodeler | TSG | Down |
| CDKN1A | Cyclin dependent kinase inhibitor 1A | Oncogene, TSG | Up |
| DDIT3 | DNA damage inducible transcript 3 | Oncogene, fusion | Up |
| ERBB2 | Erb-b2 receptor tyrosine kinase 2 | Oncogene, fusion | Down |
| FANCD2 | FA complementation group D2 | TSG | Up |

**Supplementary Table S7: COSMIC genes coordinately deregulated by IGF-1.** The Catalogue

of Somatic Mutations in Cancer (COSMIC) Cancer Gene Census lists genes implicated as cancer drivers. These included 5 genes deregulated in all 3 prostate cancer cell lines. Two genes were downregulated by IGF-1: ERBB2/HER2, possibly reflecting HER2:IGF axis crosstalk, and ATRX, a tumor suppressor gene (TSG) which is frequently lost in glioma and whose loss associates with aggressive features in osteosarcoma (14, 15). The 3 upregulated genes included *CDKN1A,* a TSG encoding cell cycle inhibitor p21 reported to induce chemoresistance in prostate cancer cells (16). Multiple functions are also reported for DNA Damage Inducible Transcript 3 (*DDIT3)* and *FADCD2. DDIT3* is a transcription factor that induces cell cycle arrest and apoptosis due to ER stress, but also promotes cancer cell proliferation, 3D growth and stemness and as a *FUS* gene fusion partner drives IGF-2 over-expression in myxoid liposarcomas (17, 18). Finally, *FANCD2* is required for inter-strand cross-link repair and recruitment of proteins required for homologous recombination, interacts with ATRX to promote replication fork restart, and its loss causes chromosome instability (19). Loss of function *FANCD2* mutation was recently identified in a patient with early-onset/familial prostate cancer (20).

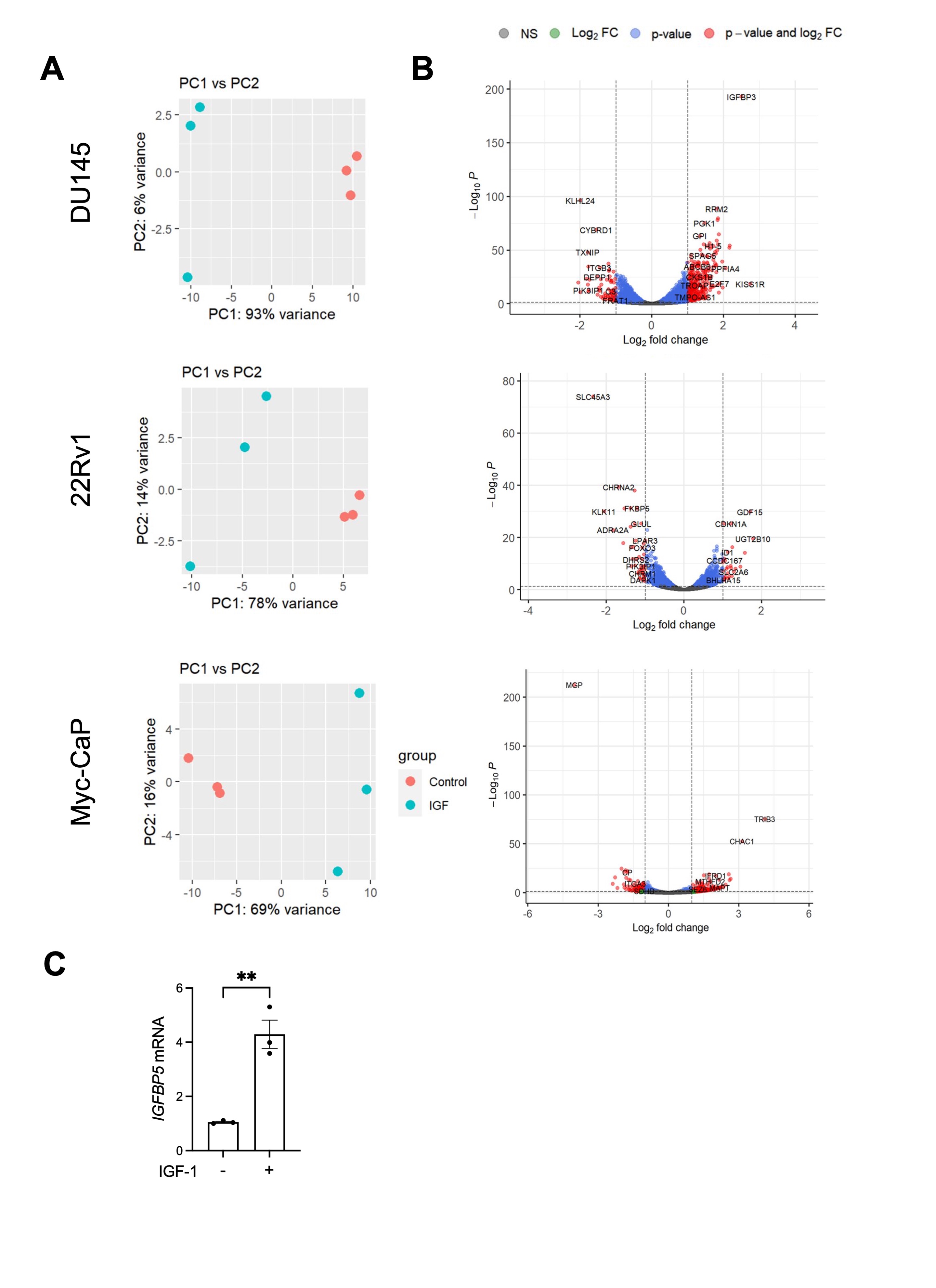

**Supplementary Figure S1. Effect of IGF-1 on transcriptome of PCa cells.** DU145, 22Rv1 and Myc-CaP cells were serum starved for 24 hrs, treated with 30 nM IGF-1 or solvent (control) for 24 hrs and RNA was extracted for sequencing. **A.** Principal component analysis (PCA) of RNA sequencing data (top 500 genes) for the three cell lines displaying the variation of each sample with respect to their transcriptomic profile. The x-axis shows the first principal component (PC1), accounting for the largest amount of variation in the experiment and the y-axis shows the second principal component (PC2). **B.** Volcano plots displaying the top high and low differentially expressed genes.

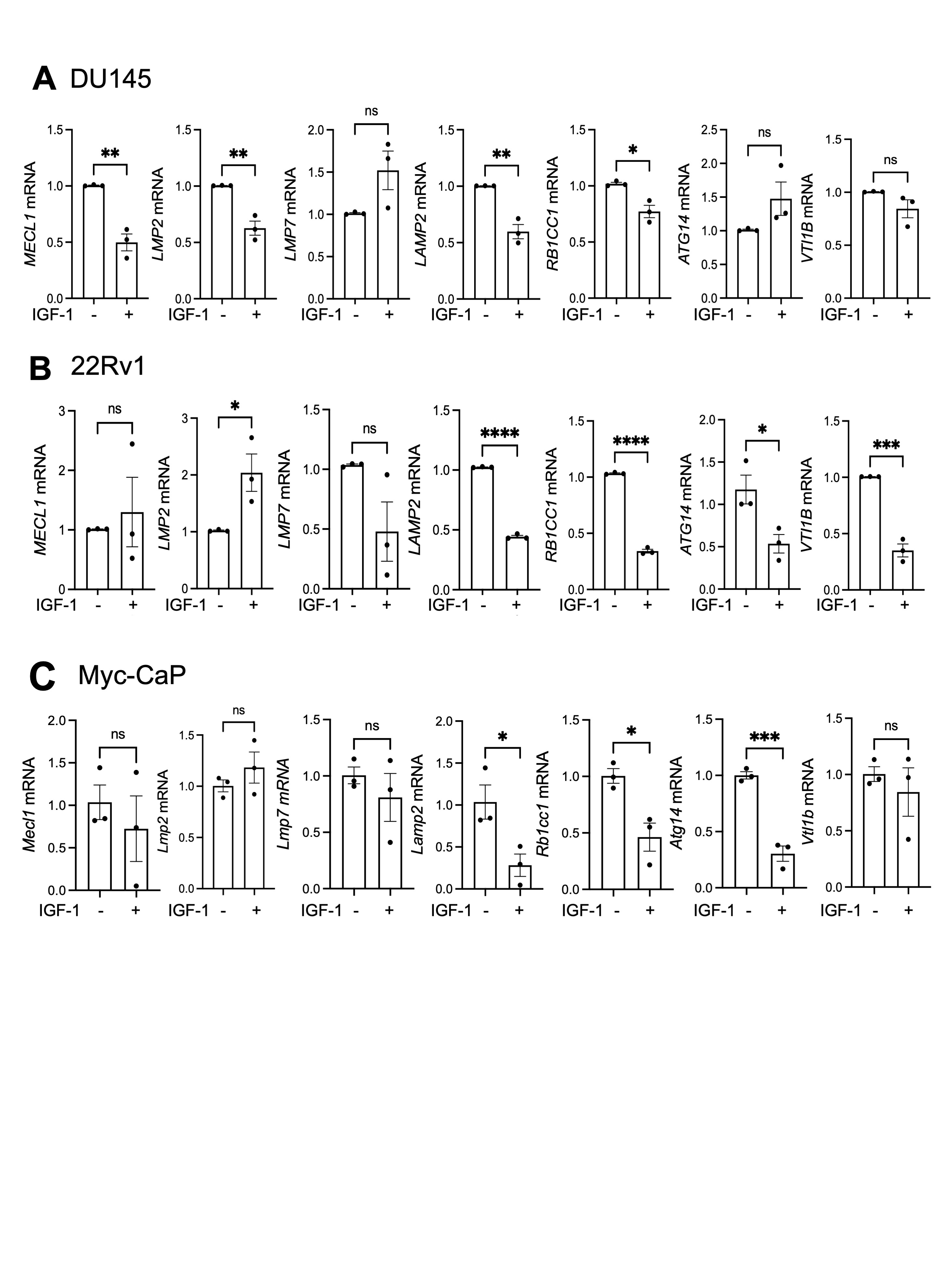

**Supplementary Figure S2. IGF-deregulated genes that influence protein degradation.** PCa cells **A.** DU145, **B.** 22Rv1 and **C.** Myc-CaP were treated for 24 hr with 30 nM IGF-1 or solvent (control) and tested for expression of the indicated genes by RT-qPCR. Graphs: mean ± SEM of three independent analyses. *p<0.05; **p<0.01; ***p<0.001; ****p<0.0001; ns, nonsignificant.

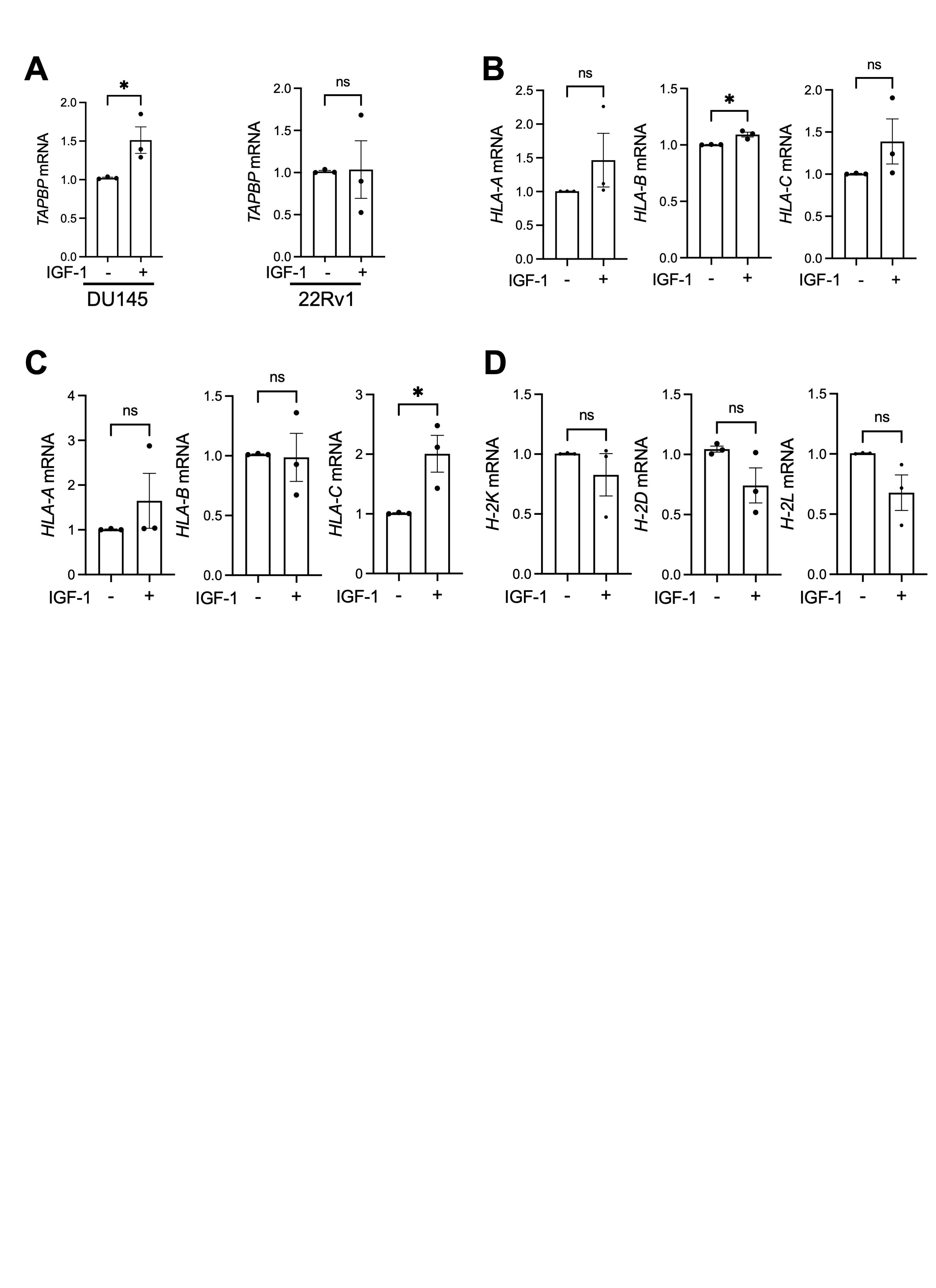

**Supplementary Figure S3. IGF-1 does not cause consistent deregulation of Class I genes in human or murine PCa cells.** PCa cells were serum starved for 24 hrs and treated with solvent (control) or 30 nM IGF-1. **A,** Effects of IGF-1 on *TAPBP* mRNA expression on left, DU145; right, 22Rv1. **B-D**. Effects of IGF-1 on expression of Class I alleles in: B, DU145; C, 22Rv1; D, Myc-CaP. Graphs represent mean ± SEM of three independent analyses (*p<0.05; ns, nonsignificant).

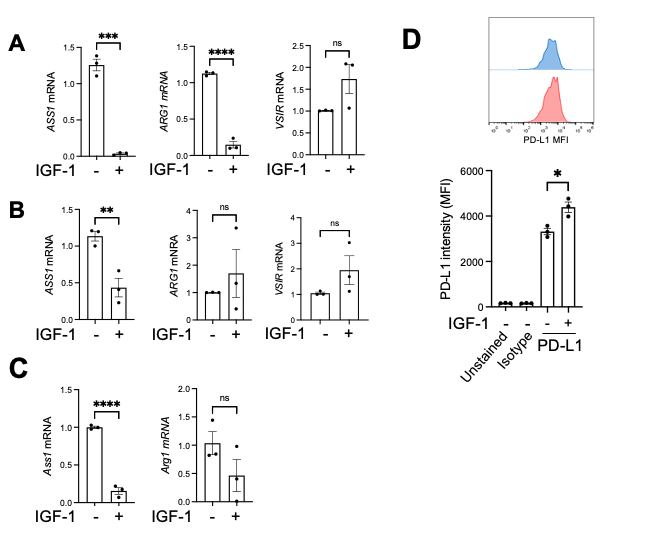

**Supplementary Figure S4. Effects of IGF-1 on expression of immune suppressors and checkpoints.** Prostate cancer cells were serum starved for 24 hrs and treated with solvent (control) or 30 nM IGF-1**. A-C.** Effects of IGF-1 on *ASS1*, *ARG1* and *VSIR* expression assessed by qRT-PCR in DU145 (A), 22Rv1 (B) and Myc-CaP (C). Vsir mRNA expression was undetectable in Myc-CaP. **D.** DU145 cells were treated with 30 nM IGF-1 or solvent (control) in full medium (10% FBS) and PD-L1 surface expression measured using flow cytometry. Upper, representative histogram; lower, expression measured as MFI, mean ± SEM from 3 independent experiments. (n=3 independent samples.*p< 0.05; **p<0.01; ***, p<0.001; ****, p<0.0001; ns, nonsignificant.

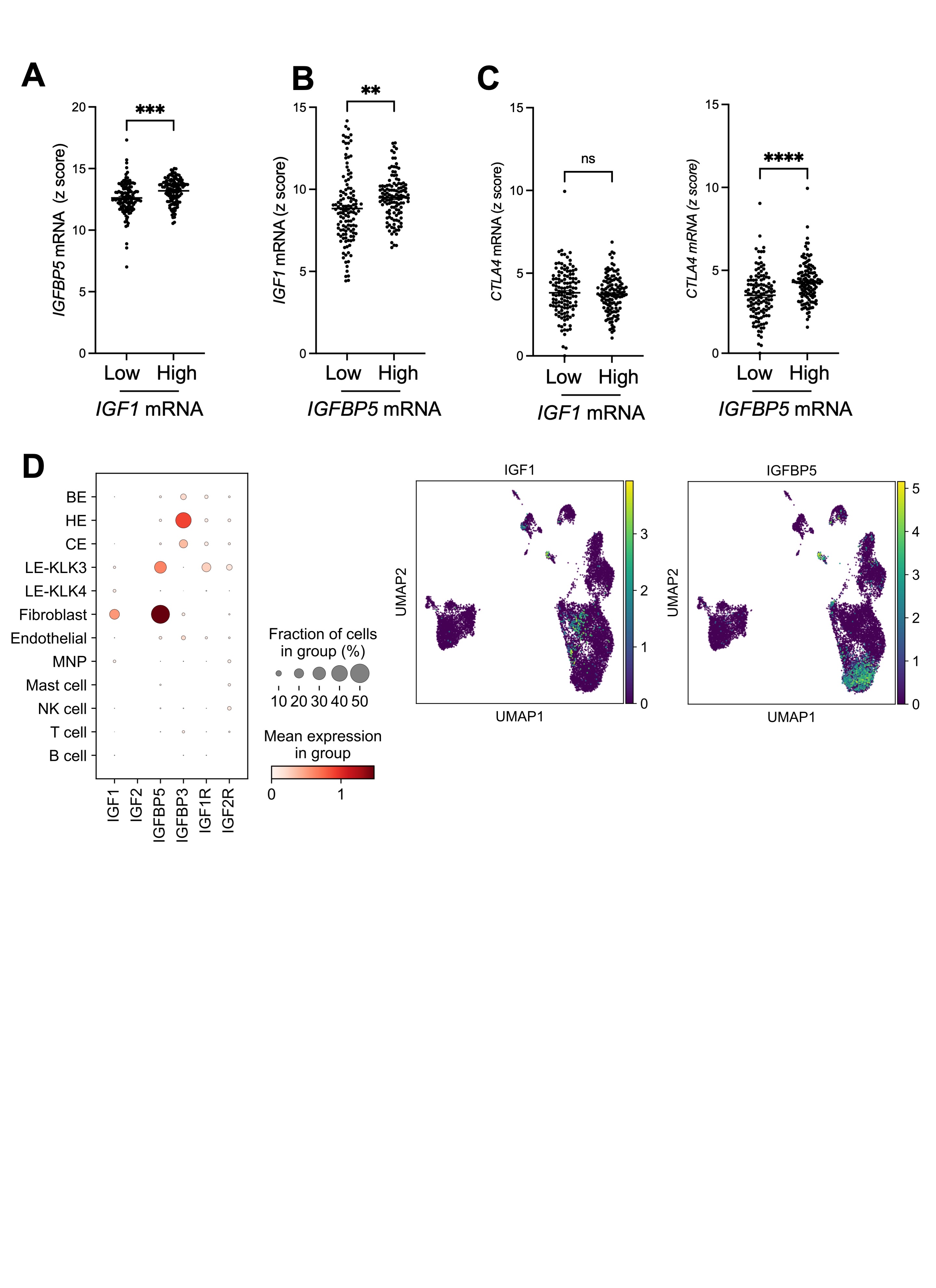

**Supplementary Figure S5. Association of IGF axis activation markers with immune checkpoints in clinical prostate cancers.** A, B and C, Tumours from TCGA prostate adenocarcinoma (n=498) were divided based on highest and lowest quartiles of tumour endogenous *IGF1* ( n= low:124, high=125) and *IGFBP5* (n= low: 124, high =125) mRNA (Z score relative to all samples). A, Association between *IGF1* mRNA and *IGFBP5* mRNA in prostate cancer tissues. B, Association between *IGFBP5* mRNA and *IGF1* mRNA in prostate cancer tissues. C, Association between *IGF1* (left) and *IGFBP5* (right) mRNA and *CTLA4* mRNA expression in prostate cancer tissue. *p<0.05; **p<0.01; ***p<0.001; ****p<0.0001; n.s., nonsignificant.

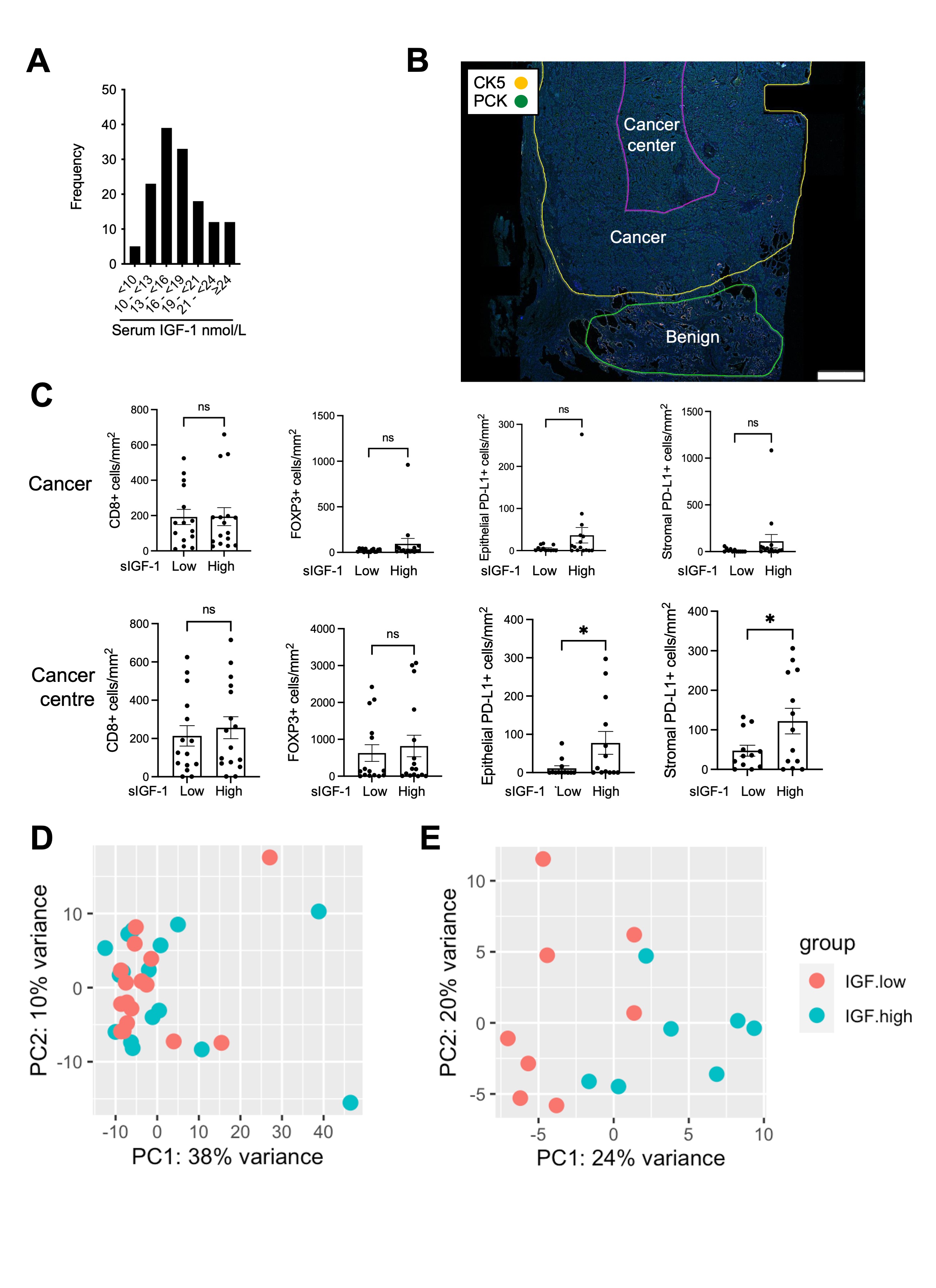

**Supplementary Figure S6. IGF axis activation associates with immune checkpoints and infiltrates in clinical PCa. A.** Serum IGF in 139 men with localized PCa recruited to ProMPT study. Assay used IDS-iSYS IGF-I assay (Immunodiagnosticsystems). All values were within normal range (7.0-31.7 nmol/L) for males >32 yr (21). **B.** PCK and CK5 from mIF of radical prostatectomy, marked up for benign, cancer, cancer center. **C.** Quantification of CD8+ T-cells, FOXP3+ Tregs, epithelial and stromal PD-L1+ cells in cancer (upper) and cancer centre (lower) in high vs low serum IGF-1 (sIGF-1) patients. Numbers in FOXP3+, CD8+ analysis: n= 15 low, 16 high sIGF-1 (1 lost due to technical issues); PD-L1+: n= 12 low, 13 high sIGF-1 patients (7 lost).
